# Tetracycline-dependent inhibition of mitoribosome protein elongation in mitochondrial disease mutant cells suppresses IRE1α to promote cell survival

**DOI:** 10.1101/2023.03.09.531795

**Authors:** Conor T. Ronayne, Christopher F. Bennett, Elizabeth A. Perry, Noa Kantorovic, Pere Puigserver

**Affiliations:** Department of Cancer Biology, Dana-Farber Cancer Institute, Boston, MA 02115; Department of Cell Biology, Harvard Medical School, Boston, MA 02115

**Keywords:** Mitochondrial disease, Tetracyclines, MALSU1, mitoribosome, IRE1α

## Abstract

Mitochondrial diseases are a group of disorders defined by defects in oxidative phosphorylation caused by nuclear- or mitochondrial-encoded gene mutations. A main cellular phenotype of mitochondrial disease mutations are redox imbalances and inflammatory signaling underlying pathogenic signatures of these patients. Depending on the type of mitochondrial mutation, certain mechanisms can efficiently rescue cell death vulnerability. One method is the inhibition of mitochondrial translation elongation using tetracyclines, potent suppressors of cell death in mitochondrial disease mutant cells. However, the mechanisms whereby tetracyclines promote cell survival are unknown. Here, we show that in mitochondrial mutant disease cells, tetracycline-mediated inhibition of mitochondrial ribosome (mitoribosome) elongation promotes survival through suppression of the ER stress IRE1α protein. Tetracyclines increased levels of the splitting factor MALSU1 (Mitochondrial Assembly of Ribosomal Large Subunit 1) at the mitochondria with recruitment to the mitoribosome large subunit. MALSU1, but not other quality control factors, was required for tetracycline-induced cell survival in mitochondrial disease mutant cells during glucose starvation. In these cells, nutrient stress induced cell death through IRE1α activation associated with a strong protein loading in the ER lumen. Notably, tetracyclines rescued cell death through suppression of IRE1α oligomerization and activity. Consistent with MALSU1 requirement, MALSU1 deficient mitochondrial mutant cells were sensitive to glucose-deprivation and exhibited increased ER stress and activation of IRE1α that was not reversed by tetracyclines. These studies show that inhibition of mitoribosome elongation signals to the ER to promote survival, establishing a new interorganelle communication between the mitoribosome and ER with implications in basic mechanisms of cell survival and treatment of mitochondrial diseases.

## Introduction

Mitochondrial diseases (MDs) are a rare, clinically heterogenous, and heritable class of diseases characterized by dysfunctional mitochondria and bioenergetic defects^1–3^. Mitochondria rely on proteins originating from both nuclear and mitochondrial genomes where mutations in genes of nuclear or mitochondrial DNA (mtDNA) origin can manifest these diseases^1–6^. The hallmark phenotype of MD includes bioenergetic defects resulting in cellular redox imbalance, inflammatory signaling, and tissue damage. MD presents in tissues of high energetic (aerobic) demands commonly resulting in neuromuscular deficiencies^2^. Specifically, mtDNA mutations in mitochondrial complex I (ex. A3796G ND1) and tRNA leucine (ex. A3243G tRNA^Leu^(UUR)) constitute the most clinically prevalent MDs, presenting as Leber hereditary optic neuropathy (LHON) and mitochondrial encephalomyopathy with lactic acidosis and stroke-like episodes (MELAS), respectively^2^. Although the precise mutations have been defined, pathological signaling mechanisms are completely unknown, and disease modifying therapeutics have not yet been realized. We have shown using a high throughput chemical screen that antibiotic tetracyclines promote survival and fitness in models of MD^7^. Antibiotics that inhibit the bacterial ribosome, like tetracyclines, also target the evolutionarily conserved mitochondrial ribosome (mitoribosome), constituting a novel treatment paradigm by modulating mitochondrial protein synthesis for MD therapy. Treatment of MD cells with tetracyclines rescues cell death from nutrient deprivation by reversing inflammatory gene expression and restoring redox homeostasis, most notably NADPH/NADP+ ratios consistent with p38 mitogen activated protein kinase (MAPK) inhibition^7^. This is also consistent with and substantiated by known mechanisms of cell death in mitochondrial complex I disease^8^. Further, tetracyclines improved survival and fitness of NDUFS4-/- mice, where the neuromuscular decline accompanying this MD was restored and was associated with attenuated neuroinflammatory signaling and suppression of hyperactivated microglia and astrocytes^7^. Interestingly, pharmacological inhibition of the endoplasmic reticulum (ER)-resident unfolded protein response (UPR) stress sensor inositol-requiring enzyme 1 (IRE1α) or downstream p38 MAPK inflammatory response can rescue MDs from nutrient deprivation^9^. These studies implicate the UPR in mediating MD cell death phenotypes under nutrient stress conditions where protein misfolding and ER stress responses are exacerbated^10, 11^. The role of mitoribosomes in signaling to the ER and specifically how tetracyclines may modulate this response to this point is unknown, prompting the current studies.

Originating from an endosymbiotic event between a eukaryotic progenitor and an α-proteobacterium, mitochondria once functioned as independent bacterial organisms. Throughout evolution, mitochondria have maintained their own genome along with transcription and translation machinery^6, 12^. Specifically, the mitoribosomes are responsible for the expression of protein subunits of the electron transport chain of mtDNA origin where translation works analogous to bacteria with functionally conserved phases of translation initiation, elongation, and termination^6^. Antibiotics that target the bacterial ribosome, like tetracyclines, are also known to target the mitoribosome^13, 14^. Specifically, tetracyclines target the bacterial ribosome by binding to 16s rRNA disrupting the accommodation of amino-acyl tRNAs in the A-site of the ribosome-mRNA complex and thereby inhibiting elongation of nascent peptides^13^. Although not yet structurally determined, it can be reasoned that tetracyclines work in a similar fashion at the mitoribosome. Mitochondrial translation is initiated upon recognition and accommodation of formyl-methionine tRNA, where subsequent amino-acyl tRNAs are recruited and translocated in a GTP-dependent fashion with a series of mitochondrial elongation factors^6^. Disruption in elongation is accompanied by the recruitment of rescue factors to maintain mitoribosome integrity and translation fidelity. Elongational stalling induced by aberrant tRNA accommodation results in a “splitting” event, where the large and small mitoribosome subunits (mtLSU and mtSSU, respectively) are dissociated and bound by a series of quality control proteins^6, 15^. First, a protein module composed of MALSU1, LOR8F8, and mtACP (MALSU1 module) binds to the mtLSU, sterically inhibiting reassociation of the mtSSU^15^. Subsequently a series of rescue factors (MTRES1 and mtRF-R) bind and stabilize nascent peptide and P- site tRNA^15^. These quality control proteins identify a means by which the cell can recycle native-disrupted or inhibited mitoribosomes to bypass energetically and synthetically costly rounds of ribosome biogenesis. The role of these proteins, and largely of the mitoribosomes, in signaling cell fate decisions has not yet been described, offering a starting point for the current work toward the investigation into the initial molecular mechanisms of tetracycline-induced cell survival.

In the current work we investigate signaling mechanisms that originate at the mitoribosome in response to tetracyclines in MD. Here, we illustrate that tetracyclines mediate the recruitment of mitochondrial quality control protein MALSU1 to the mtLSU, indicating that elongational stalling and splitting are initial responses to tetracycline treatment. Using CRISPR-Cas9 technology, we identify that MALSU1, but not other rescue factors, is required for tetracycline-induced survival response in MD cells under nutrient deprived conditions. We further show that tetracyclines reverse cell-death signaling in the ER through the attenuation of IRE1α oligomerization and inhibition of effector UPR function. MALSU1 deficient MD cells sensitive to nutrient stress exhibit heightened sensitivity toward the activation of ER stress, which is not reversed by tetracyclines. This work highlights a novel signaling between the mitochondria and ER that can be leveraged for the treatment of MD.

## Results

### Tetracyclines require mitoribosome splitting and quality control protein MALSU1 to induce survival in ND1 complex I mutant cells

Mutations in mitochondrial genes result in sensitivity to nutrient deprivation where glucose deprivation can induce cell death. Specifically, cells cultured in equimolar concentrations of galactose in place of glucose become dependent on mitochondrial metabolism, where mitochondrial mutations render these cells particularly sensitive to cell death^7–9, 16^. We previously illustrated that inhibition of mitochondrial translation by tetracyclines rescues mitochondrial disease models from cell death by a currently unknown mechanism^7^. Here, utilizing mitochondrial disease complex I (A3796G ND1) mutant cybrid U2OS cells, we interrogate mitoribosome quality control pathways as potential mediators of tetracycline-induced rescue under glucose deprivation.

Recent cryogenic electron microscopy studies in phosphodiesterase 12 (PDE12) mutant HEK293 cells indicate that mitoribosome elongational stalling induced by aberrant tRNA accommodation promotes mitoribosome splitting and recruitment of quality control factors^15^. Doxycycline inhibits bacterial protein synthesis by binding to the bacterial SSU rRNA in the decoding region, stalling translation by disrupting the accommodation of tRNA^13^. This mechanism is similar to defective tRNA processing in PDE12 mutant cells^15^. In this regard, we sought to investigate mitoribosome elongational stalling and quality control factors as initial molecular signals in promoting doxycycline-induced survival in ND1 mitochondrial mutant cells. ND1 mutations are commonly found in MD patients^2^, and we have previously used this cell line model to validate genes and compounds, including tetracyclines, that rescue cell death under glucose-deprivation conditions^7–9^. The mitochondrial translation cycle includes distinct stages of initiation, elongation, and termination, where conserved classes of proteins regulate progression between each phase^6^. To investigate if active mitochondrial elongation is required for survival in ND1 mutants, we compared the effects of elongation-inhibition of doxycycline with a distinct inhibitor, N-formyl methionine mimetic actinonin, that stalls mitochondrial translation by inhibiting peptide deformylase^17, 18^ (PDF, **Fig. 1A**). PDF removes N-formyl groups on nascent peptides as they emerge from the ribosome exit tunnel prior to folding^17, 18^, and its mechanism of inhibition of mitochondrial translation is distinct to that of tetracyclines. These studies revealed that actinonin did not promote survival in ND1 cells at a wide range of concentrations (**Fig. 1A**). We have previously identified^7^ that other elongation inhibitors pentamidine^19^ and retapamulin^13, 20^ promoted survival similar to doxycycline, substantiating this mechanism. These results suggest that general suppression of mitochondrial protein synthesis is not sufficient, but rather a signal must be initiated at the elongating mitoribosome, to promote survival.

**Figure 1.**
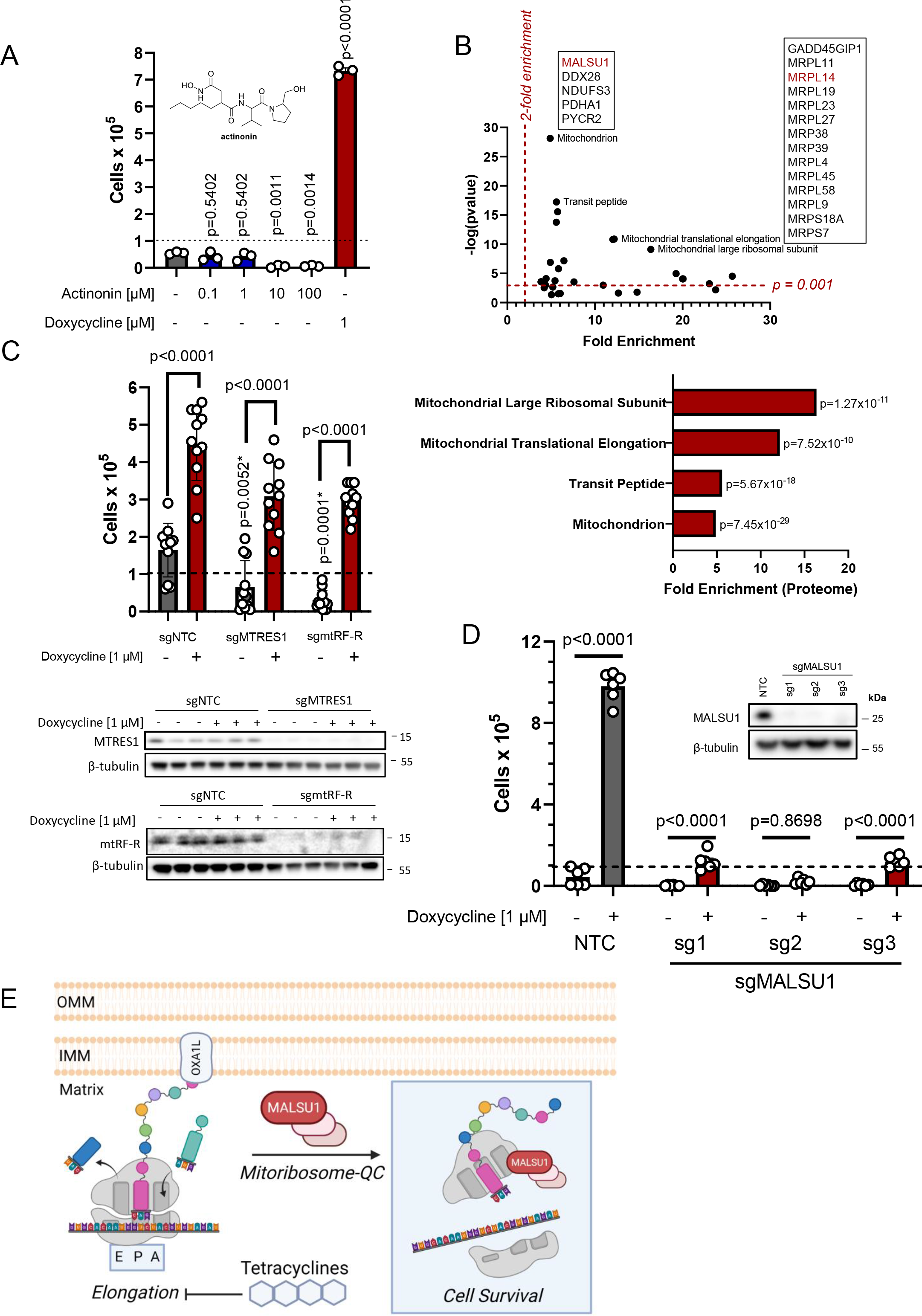
Tetracyclines require mitoribosome splitting and quality control protein MALSU1 to induce survival in ND1 mutant cells. Mitoribosome elongational stalling recruits’ quality control factors^15^. (**A**) ND1 cells treated with a range of concentrations (0.1, 1.0, 10, 100 µM) of peptide deformylase inhibitor actinonin do not promote survival under galactose conditions (n=3 independent biological samples). Doxycycline (1 µM) potently rescued ND1 cells. (**B**) DepMap drug-repurposing proteomics database indicates enrichment of mitochondrial and more specifically mitoribosome proteins in the context of doxycycline. Gene-ontology enrichment was determined using DAVID pathway analysis, where significant proteins were determined (p<0.001)^41^ and compared to entire proteome. Note enrichment of MALSU1 associated with surviving cells in the context of doxycycline. (**C**) Depletion (CRISPR-Cas9) of mitochondrial quality control rescue factors MTRES1 and mtRF-R did not reverse capacity of tetracyclines to rescue ND1 cells from galactose induced cell death (n=11 biological replicates over N=4 experiments). P-values* with asterisk denote comparisons with DMSO treated NTC cells. (**D**) MALSU1 expression is required for tetracyclines induced survival. During survival experiments, dashed-line represents ND1 seeding density, where cells were exposed to glucose deprivation (galactose supplementation) for 4 days (n=6 biological replicates over N=2 experiments). (**E)** Proposed model illustrating doxycycline-induced and MALSU1-dependent cell survival based on activation of mitoribosome quality control (QC)^15^. Cell survival responses to tetracycline treatment depends on MALSU1 but not MTRES1 or mtRF-R. Created with BioRender.com.

In an unbiased approach to identify factors involved in doxycycline rescue, we utilized the Dependency Map (DepMap) proteomics datasets where repurposing screens have been performed to investigate the utility of current FDA approved small molecules, including doxycycline, for cancer treatment^21, 22^ (**Fig. 1B**). Here, positive correlation between the proteome and doxycycline identified proteins involved in cell death. Hence, we sought to classify proteins that negatively correlate with doxycycline, promoting survival in line with its mechanism in models of MD^7^. Gene-ontology enrichment analysis identified high (negative) correlation with numerous mitochondrial protein sub-classes “mitochondrion” and “transit peptide”, and more specifically the mitoribosome including “translational elongation” and “large mitoribosome subunit” (**Fig. 1B**). Interestingly, these studies identified survival correlation with mitoribosome quality control protein MALSU1, in line with potential elongational stalling and splitting mechanism described above. To note, MTRES1 and mtRF-R did not correlate with doxycycline treatment. MRPL14, an accessory subunit of the mtLSU that MALSU1 binds during splitting^15, 23^, was also identified in this screen, further substantiating the importance of this binding module in surviving cancer cells and potentially in mitochondrial mutant cells in the context of doxycycline mechanisms of action (**Fig. 1B**).

To interrogate the mitoribosome quality control pathway in facilitating tetracycline survival mechanisms in mitochondrial ND1 mutant cells, we utilized CRISPR Cas9 gene editing to knock-out (KO) respective proteins involved in this pathway. Elongational stalling results first in splitting of the large and small subunit where MALSU1, along with LOR8F8 and NDUFAB1 (MALSU1 module), binds and stabilizes the large subunit. Rescue factors MTRES1 and mtRF-R then bind and stabilize the nascent peptide and tRNA. We first knocked out MTRES1, mtRF-R, and MALSU1 to identify which, if any, proteins may be involved in doxycycline survival (**Fig. 1C-D**). These studies illustrated that doxycycline was still able to rescue ND1 cell death in the absence of MTRES1 or mtRF-R (**Fig. 1C**), but survival was significantly reduced in the absence of MALSU1, and thereby un-bound LOR8F8 and NDUFAB1 (**Fig. 1D**). It was also noted that basal survival rates of ND1 cells in galactose were drastically reduced in MTRES1 and mtRF-R KO conditions (**Fig. 1C**), indicating that these cells rely on mitoribosome quality control under nutrient stress, but doxycycline specifically works through the MALSU1 module to promote survival (**Fig. D- E**).

### Tetracyclines mediate MALSU1 recruitment to the mitoribosome large subunit

To investigate the mechanisms whereby MALSU1 was required for mitochondrial mutant disease cell survival under nutrient stress, we sought to further investigate doxycycline-dependent regulation of MALSU1 binding to the mitoribosome. To study these interactions at the purified mitoribosome, we utilized HEK293 suspension cells (Expi293F), a model previously used to study mitoribosome elongational stalling and quality control^15^. ND1 mutant cybrid cells grown in monolayer culture did not provide sufficient mitochondria to purify appreciable yields of the mitoribosome and thus, Expi293F suspension cells were used as an initial cellular model system to study potential changes in the molecular composition and architecture of the doxycycline-inhibited mitoribosome. Rapid isolation of the mitoribosome was achieved through successive sucrose gradient centrifugations to obtain first the mitochondria, which were then solubilized in dodecyl-β- D-maltoside (β-DDM) and cardiolipin. This efficiently extracted mitoribosomes from the inner mitochondrial membrane for fractionation of the mtSSU, mtLSU, and monosomes^15^. Expi293F cells were treated with doxycycline for 48 hours to induce elongational stalling and then mitoribosomes were isolated and sedimented to compare with vehicle-treated cultures. Doxycycline-treated mitochondria, and more drastically the mitoribosomes, exhibited increased MALSU1 protein content (**Fig. 2A**). These results are consistent with increases in MALSU1 binding to the mtLSU as previously noted (**Fig. 1E**) and may reflect increases in its mitochondrial import and/or stability in response to doxycycline. To investigate whether MALSU1 was enriched at the large subunit in doxycycline-treated cultures, we sucrose-gradient-fractionated mitoribosomes to separate and analyze the location of MALSU1. Here, we found that MALSU1 was enriched at the mtLSU upon doxycycline treatment (**Fig. 2B**). Additionally, we observed a shift in the sedimentation of the MALSU1-bound mtLSU with doxycycline treatment, indicating a potential change in the size and/or molecular shape of the large subunit in this condition when compared to controls (**Fig. 2B**). To validate these effects in ND1 cybrid cells, crude mitochondria were isolated, solubilized with β-DDM and cardiolipin, and fractionated by sucrose gradients in a similar fashion as described above. Additionally, to investigate the necessity of MALSU1 in shifting the sedimentation pattern of the doxycycline treated large subunit, mitochondria from CRISPR-Cas9 MALSU1 depleted ND1 cells were analyzed in parallel. As observed in Expi293F suspension cells, doxycycline treatment in ND1 cells resulted in an increase in mitochondrial MALSU1 (**Fig. 2C**) that was enriched at the large subunit (**Fig. 2D-E**). The shift in large subunit sedimentation was again observed with doxycycline treatment (**Fig. 2D**) mimicking Expi293F cultures (**Fig. 2B**). Sucrose gradients of MALSU1 depleted mitochondria illustrate that the doxycycline treated mitoribosome sedimentation is again altered (**Fig. 2F**), indicating that doxycycline may split the large subunit independent of MALSU1, but MALSU1 binding to the large subunit is required for survival (**Fig. 1D-E**).

**Figure 2.**
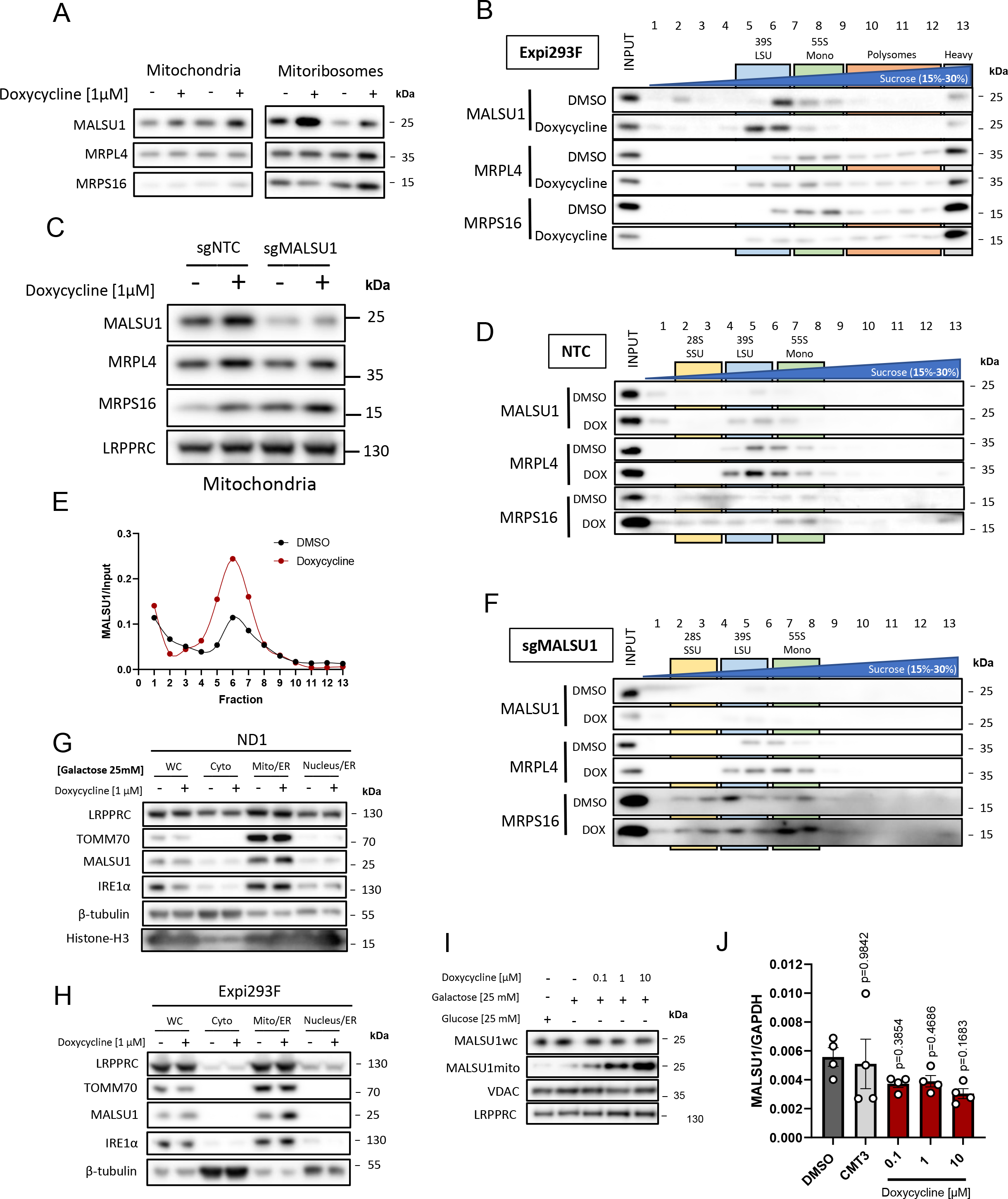
Tetracyclines mediate MALSU1 recruitment to the mitoribosome large subunit. (**A**) Experimental scheme for the isolation and sedimentation of mitoribosomes from Expi293F suspension cells. Treatment of isolated mitochondria with Dodecyl-beta-D- maltoside (β-DDM) and cardiolipin efficiently extracts the mitoribosome from the inner mitochondrial membrane for sucrose gradient sedimentation. (**B**) Isolated mitochondria and mitoribosomes from Expi293F exhibit enrichment of MALSU1 in doxycycline treated cultures, that is (**C**) associated with the large subunit of the mitoribosomes evidenced by sucrose gradient sedimentation (15-30% sucrose). Sedimentation of large and small subunit were assessed by MRPL4 and MRPS16, respectively. Duplicates of mitochondria and mitoribosomes in (**B**, independent experiments) are the inputs for (**C**). Note shift in MALSU1 and MRPL4 upon doxycycline treatment. (**D**) Isolation of crude mitochondria from doxycycline (1µM) treated ND1 cells (non-target control (NTC) and sgMALSU1). Note increased MALSU1 expression in doxycycline-treated mitochondria. (**E**) Crude mitochondria from ND1-NTC cells were sucrose gradient (15-30%) sedimented and analyzed for MALSU1 as in (**C**). Note shift and enrichment of MALSU1 and MRPL4 with doxycycline treatment. (**F**) Quantitation of MALSU1 elution profile in sucrose gradients of crude mitochondria from ND1 cells derived from (**E**), normalized to input signal of MALSU1. (**G**) Sucrose gradient (15-30%) sedimentation of MALSU1 depleted ND1 cells with or without doxycycline treatment. Crude mitochondria from (**D**) are inputs for (**E**) and (**G**). (**H**-**I**) Cell fractionation of (**H**) ND1 and (**I**) Expi293F suspension cultures treated with doxycycline. Note enrichment of MALSU1 specifically in mitochondrial fractions. (MAM; mitochondrial associated ER membrane). (**J**) ND1 cells treated with increasing concentrations of doxycycline illustrate dose-dependent increases in MALSU1 specifically at the mitochondria without altered whole cell (WC) expression levels or changes in total mitochondria (VDAC, LRPPRC). (**K**) Quantitative PCR of ND1 cells (galactose, 48 hr) treated with varying concentrations of doxycycline or inactive tetracycline analog CMT3 indicate mRNA expression levels of MALSU1 are unchanged (n=4 biological replicates across N=2 independent experiments).

To further investigate the sub-cellular localization of MALSU1 in response to doxycycline, we fractionated ND1 (**Fig. 2G**) and Expi293F (**Fig. 2H**) cells to identify if MALSU1 is enriched in specific compartments in response to elongational stalling. These experiments illustrated that overall levels of MALSU1 (whole cell) are unchanged in response to doxycycline treatment, but it is specifically enriched at the mitochondria without changes in mitochondrial matrix protein LRPPRC or outer-mitochondrial membrane TOM70 (**Fig. 2G-H**). Additionally, dose-dependent increases in mitochondrial localization without changes whole cell protein (**Fig. 2I**) or transcript levels (**Fig. 2J**) further support that regulation of MALSU1 at the mitochondria is post-translational. These studies support the notion that MALSU1 binding to the mitoribosome large subunit, and not simply its increased expression, is the molecular action driving tetracycline-induced survival responses and illustrate a direct association between the extent of elongation inhibition and MALSU1 recruitment.

Taken together, elongational stalling caused by doxycycline results in mitoribosome splitting where doxycycline acts as a “splitting factor” and MALSU1 binds and stabilizes the large subunit to provide a pro-survival signal. Doxycycline efficiently rescues ND1 mutant cells lacking the downstream rescue factors MTRES1 and mtRF-R (**Fig. 1C**) and thus, when elongation is stalled doxycycline-dependent MALSU1 binding provides a pool of ribosomes that promote survival under nutrient stress.

### Tetracyclines promote survival through MALSU1-dependent suppression of ER stress IRE1α signaling

We have previously reported that pharmacological inhibition of ER stress sensor/transducer IRE1α promotes survival in mitochondrial complex I disease cells^9^, implicating ER stress in promoting cell death in mitochondrial defective respiration. To determine whether doxycycline-induced survival impinges on this pathway, we tested if doxycycline treatment reversed the activation of X-box binding protein 1 short fragment (XBP1s), an ER-stress-induced IRE1α cleavage product that indicates activation of the unfolded protein response (UPR)^10, 11^. Western blot analysis indicated that XBP1s is strongly and specifically activated in ND1 cells, but not WT cells, after 48 and 72 hours under glucose restriction conditions (**Fig. 3A**). Doxycycline treatment efficiently reduced activation at 48 hours and completely reversed at 72 hours consistent with cell survival (**Fig. 3A**), implicating doxycycline in resolving nutrient-dependent ER stress. Interestingly, CMT3, a tetracycline analog that does not exhibit mitoribosome protein translation inhibition or rescue mitochondrial mutants^7^, does not reverse the activation of XBP1s (**Fig. 3A**). These studies indicate that inhibition of mitochondrial translation elongation is required for the pro-survival resolution of ER stress. These effects were extended to additional markers of ER stress including dose-dependent decreases in ER lumen UPR chaperone BiP and downstream p38 MAPK phosphorylation in a similar fashion to XBP1s (**Fig. 3B**). In fact, we have previously illustrated that pharmacological inhibition of p38 (SB203580) rescues survival in ND1 mutant cells concomitant with metabolic changes similar to that of doxycycline treatment implicating MAPK signaling in the survival mechanism^7^. Treatment of ND1 cells with IRE1α inhibitor 4µ8C reverses the galactose- induced phosphorylation of p38 (**Fig. S1A**) similar to doxycycline or chemically distinct and active pentacycline-based mitoribosome inhibitor 7002 but not CMT3 (**Fig. S1B**). This illustrates that MAPK activation is downstream of IRE1α and can be controlled by mitoribosome inhibition under these conditions. Unfolded proteins in the ER bind to IRE1α on its luminal domain promoting dimer/oligomerization which activates the cytosolic kinase and RNAase domains, transducing ER stress^11^. To further investigate doxycycline effects on IRE1α activation, we analyzed its oligomerization state using blue-native polyacrylamide gel electrophoresis (BN-PAGE, **Fig. 3C**) as previously described^24^. In whole cell extracts, we did not observe changes in the oligomerization state with doxycycline treatment, inconsistent with results on XBP1s activation. It has been previously reported that IRE1α can be enriched at mitochondrial ER contact sites^25^, a potential sub-cellular location wherein the oligomerization state is regulated. With doxycycline mode of action at the mitoribosome, we isolated crude mitochondria containing ER fractions from ND1 cells and performed BN-PAGE to analyze the IRE1α oligomerization status in the ER concentrated at the mitochondria. Here, we observed a marked dose-dependent decrease in oligomerization state of IRE1α with doxycycline treatment without changes in total IRE1α levels or associated mitochondrial markers (**Fig. 3C**). These results indicate that ND1 cell-survival decisions through IRE1α are to some extent controlled at the mitochondria-associated ER, and that a signal may be initiated at the mitochondrial ribosome upon elongational stalling with doxycycline treatment.

**Figure 3.**
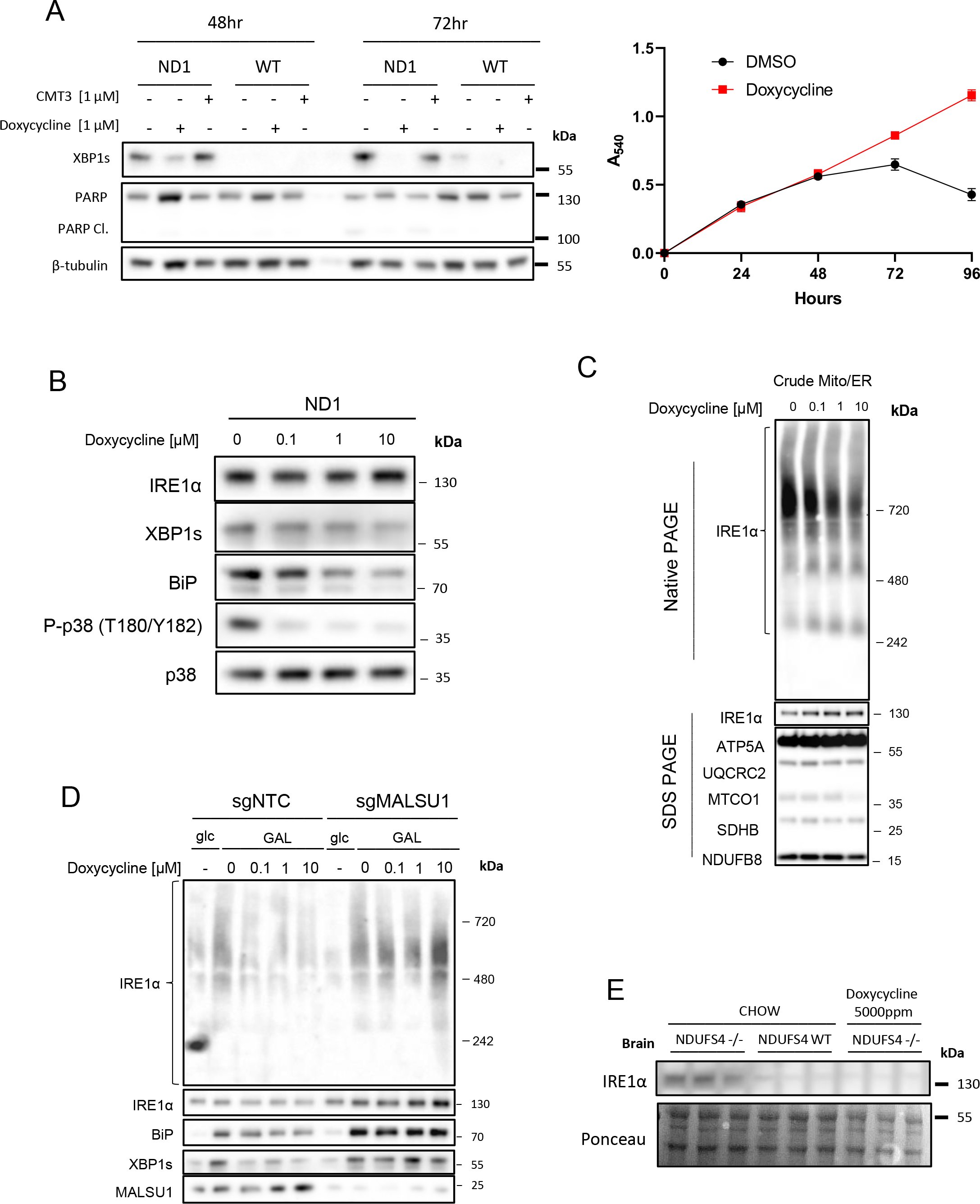
Tetracyclines promote survival through MALSU1-dependent suppression of ER stress IRE1α signaling. (**A**) Doxycycline (1 µM) reverses activation of IRE1α as evidenced by decrease in XBP1s in ND1 cells (galactose, 48 and 72 hours). Inactive CMT3 tetracycline analog does not reverse XBP1s. Suppression of XBP1s at 72 hours is associated with reversed PARP cleavage, consistent increased survival at later time points. Note galactose does not induce XBP1s in WT cultures. Survival curve over 96 hours indicates cell death initiates at the 48 hour time point in vehicle treated cells (DMSO, points represent the average A540 signal ± SD (n=2 biological replicates). (**B**) Doxycycline dose-dependently inhibits activation of XBP1s, ER chaperone BiP, and phosphorylation of p38 MAPK in ND1 cells (galactose, 48 hr). (**C**) Blue-Native PAGE indicates dose- dependent reductions in high-order IRE1α oligomers in isolated crude mitochondria (Mitochondria-ER). (**D**) IRE1α oligomerization status in mitochondria-ER fractions and effector UPR function in ND1 cells is MALSU1 dependent. Note increased oligomerization of IRE1α upon galactose treatment that it attenuated in doxycycline treated cultures (non- target control, NTC), with reductions in BiP and XBP1s. IRE1α oligomerization status was not attenuated by doxycycline in ND1 cells lacking MALSU1, and consistent with BiP and XBP1s. (**E**) IRE1α expression levels in the brains of NDUFS4-/- mice indicate increased expression when compared to WT and doxycycline-supplemented counterparts. Results represent whole brains isolated from three independent mice under each indicated condition (n=3 independent biological replicates).

MALSU1, over other mitoribosome quality control proteins, is required for doxycycline-induced survival (**Fig. 1D&E**). To investigate if MALSU1 is required for doxycycline-dependent inactivation of IRE1α we utilized MALSU1 depleted ND1 cells in a similar analysis of oligomerization and downstream XBP1s activation when compared to control cells. In control and MALSU1 KO cells, we observe an increase in IRE1α oligomerization at mitochondrial contacts, along with XBP1s and BiP activation with galactose treatment (**Fig. 3D**). However, unlike control cells, IRE1α oligomerization, XBP1s cleavage, and BiP activation are not repressed in MALSU1 KO cells with doxycycline treatment, consistent with the survival experiments shown in Figure 1 (**Fig. 3D**). These studies show that tetracyclines require MALSU1 to modulate IRE1α-mediated UPR activity in ND1 mitochondrial mutant cells.

NDUFS4 KO mice are a widely employed pre-clinical model of mitochondrial disease, largely characterized by neuromuscular decline^26^. Previously, we used this model to investigate the utility of doxycycline *in vivo*, where we found doxycycline supplemented mice exhibited improved survival and fitness when compared to controls^7^. Specifically, doxycycline reversed neurological defects in NDUFS4 KO brains. Thus, we investigated the status of IRE1α in the brains of NDUFS4 KO mice (**Fig. 3E**). We found that NDUFS4 KO brains exhibited marked increased expression of IRE1α protein levels when compared to their control and doxycycline treated counterparts (**Fig. 3E**). These results indicate that chronic treatment of doxycycline may attenuate the UPR in brains with mitochondrial dysfunction, suppressing IRE1α. Taken together, survival signaling at elongating mitoribosmes requires MALSU1 where cross-talk between mitoribosome quality control and ER-stress effectors highlight an important survival mechanism in mitochondrial disease mutants under nutrient stress.

Canonical regulation of IRE1α includes the interplay between inhibitory-binding of BiP to monomeric IRE1α on the luminal domain and the activating-accumulation of unfolded proteins, releasing BiP to chaperone and maintain protein homeostasis^11^. This release allows for dimerization of IRE1α, transducing initial pro-survival UPR, where prolonged ER stress incapacitates BiP (and other) chaperone activities. Unfolded proteins bind to the IRE1α luminal domain directly, causing oligomerization activating cell death responses^11^. MARCH5, a mitochondrial outer membrane ubiquitin ligase, has recently been implicated in attenuating the oligomerization of IRE1α under prolonged ER stress^25^. With marked decreases in IRE1α oligomerization upon mitochondrial translation inhibition with doxycycline (**Fig. 3C**), we sought to investigate if this regulation was mediated through MARCH5. CRISPR Cas9 KO of MARCH5 sensitized ND1 cells to galactose-induced ER stress with heightened levels of XBP1s and P-p38, but this response could still be reversed by doxycycline treatment (**Fig. S2**). Importantly, doxycycline still rescued MARCH5 depleted ND1 cells from galactose induced cell death (**Fig. S2**), implicating a novel IRE1α regulatory mechanism originating at the mitochondria involving the translation machinery and MALSU1 that can be activated with elongational stalling.

### Mitochondrial ND1 mutant cells exhibit increased ER protein loading that is attenuated by tetracyclines

Intrigued by the ability of tetracyclines to suppress ER stress in mitochondrial mutants, we sought to further evaluate mechanisms of ER protein homeostasis. ER stress and homeostasis are largely regulated by (1) ER associated protein degradation (ERAD) and (2) ER protein translocation or insertion^27^. To test if tetracyclines are resolving ER stress by activating ERAD, we utilized specific inhibitor of p97 induced protein degradation eeyarestatin I^28, 29^ (ESI, **Fig. 4A-C**). It was noted in these experiments that mitochondrial mutant cells were more dependent on ERAD for survival under nutrient stress when compared to WT, as the potency of ESI was dramatically increased under galactose conditions in ND1 mutant cells (**Fig. 4A**) but unchanged in WT cells (**Fig. 4B**). Galactose-induced phosphorylation of p38 MAPK and XBP1 splicing were increased in ND1 when compared to WT cells, exacerbated by ESI, and suppressed by IRE1α inhibition, illustrating that ERAD modulates MAPK and XBP1s signaling downstream of UPR (**Fig. S1A**). However, we observed that treatment with ESI did not reverse the ability of tetracyclines to promote survival in mitochondrial mutant cells under nutrient stress (**Fig. 4C**), prompting our investigation into ER-protein translocation as a means of tetracycline rescue. To assess the effects of tetracyclines on ER protein translocation, we probed tetracycline-treated mitochondrial mutant cells and isolated microsomes (ER) for the accumulation of cathepsin-C, a glycosylated and secreted protein that can be used as a marker for translocation in ER fractions^30^. Here, we found that microsomes derived from ND1 cells exhibit a substantial increase in cathepsin-C when compared to those derived from WT cells and tetracyclines suppressed this accumulation (**Fig. 4D&E**). Expression or stability of cathepsin-C were not altered in whole cell extracts or microsomes as indicated by cycloheximide chase experiments, and expression levels were comparable between WT and ND1 cells (**Fig. 4D&E**) suggesting that ER protein loading is specifically increased upon mitochondrial mutation, consistent with increased BiP in the ER from ND1 cells that is reduced by tetracyclines (**Fig. 4E**). Due to the necessity of MALSU1 in promoting survival, we investigated the tetracycline-induced effects on ER protein loading in mitochondrial mutant cells lacking MALSU1. We found that tetracyclines were unable to modulate cathepsin-C microsome loading in MALSU1-deficient cells (**Fig. 4F**), further implicating this mitoribosome quality control response in the regulation of ER-protein loading and ER stress. ER-protein translocation and specifically the secretory pathway is initially regulated by the signal recognition particle (SRP), which binds to the ribosome nascent chain complex to promote co-translational translocation into the ER lumen^31, 32^. We observed a marked decrease in SRP subunit SRP14 in doxycycline treated microsomes from ND1 cells (**Fig. 4E**), that was not observed in the absence of MALSU1 (**Fig. 4F**) further supporting a decrease in ER-protein translocation in doxycycline-treated cultures. Enhanced ER protein loading in mitochondrial mutant cells may explain the increased dependence of cells with ND1 mutation on ERAD (**Fig. 4A**) and the enhanced sensitivity to ER stress as presented in figure 3 (**Fig. 3A**). We present a model whereby inhibition of mitoribosome elongation with tetracyclines activates mitoribosome quality control and resolves ER stress and IRE1α signaling in mitochondrial mutant cells by a currently unknown signaling mechanism (**Fig. 4G**).

**Figure 4.**
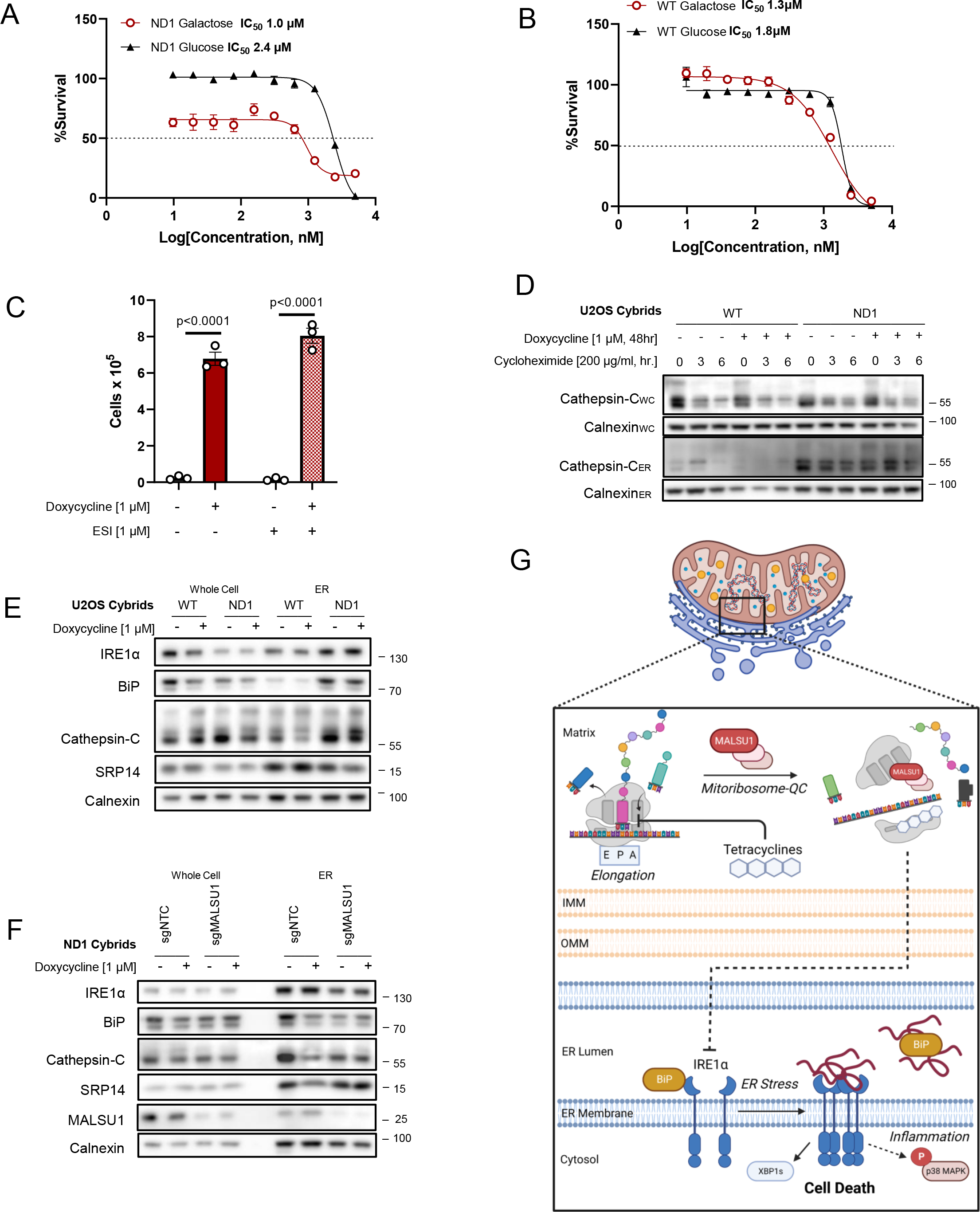
Mitochondrial mutant cells exhibit increased ER protein loading that is attenuated by tetracyclines. Dose-dependent treatment with p97 specific ERAD inhibitor eeyarestatin I (ESI) shows (**A**) mitochondrial mutant cells are more dependent on ERAD when compared to (**B**) WT as indicated by enhanced potency (IC50) of ESI in ND1 cells under galactose-stress conditions (n=2 biological replicates). (**B**) Note ESI potency is unchanged between glucose and galactose conditions in WT cells (IC50). (**C**) Inhibition of ERAD with ESI does not reverse doxycycline-induced survival (n=3 independent biological replicates). (**D**) ER protein cathepsin-C loading is increased in isolated microsomes (ER) upon mitochondrial mutation, which is attenuated by doxycycline. Whole cell (WC) expression levels are unchanged between WT and ND1 cells. Cycloheximide treatment indicates comparable half-lives between control and doxycycline treated cultures and isolated microsomes. (**E**) Microsomal cathepsin-C levels that are increased upon ND1 mutation are attenuated by doxycycline, and associated with decreases in BiP and SRP14 in these fractions. (**F**) Depletion of MALSU1 in ND1 cells reverses doxycycline regulation of BiP, cathepsin-C, and SRP14 at the microsome. (**G**) Model describing inhibition of mitochondrial translation with tetracyclines signals the suppression of ER stress and UPR activation through IRE1α. Dashed lines indicate indirect association. Created with BioRender.com

## Discussion

MD represents a clinically heterogenous and difficult to treat subset of metabolic diseases, where mechanisms of disease progression and pathological signaling are largely unknown^1–4^. Here, we present a model whereby inhibition of mitochondrial translation with tetracyclines splits the mitoribosome and recruits MALSU1, reducing lethal ER stress and IRE1α signaling (**Fig. 4G**). Mechanisms of cross-talk between the mitochondria and ER have been widely explored in normal physiology and disease processes including but not limited to autophagy, bioenergetics, calcium signaling, redox balance, protein import, and metabolite exchange^10^; aspects commonly accompanied by pathological transformation in MD. Here we show for the first time a functional cross-talk between a mitoribosome associated protein and the unfolded protein response (UPR) that expands our current knowledge and scope of interorganelle communication.

As the protein processing hub of the cell, the ER promotes the proper glycosylation, folding, membrane insertion, and delivery of proteins to their subcellular destination^27^. Defects in these processing mechanics results in accumulation of misfolded proteins in the ER lumen, activating the UPR through a series of ER-membrane stress sensors IRE1α, PERK, and ATF6^10, 27^. IRE1α, the most conserved UPR protein, contains an ER luminal domain that interacts with ER chaperones (namely BiP) and/or misfolded proteins, and a cytosolic kinase/RNAase domain responsible for transducing luminal protein stress^10, 11, 27^. Heightened levels of misfolded proteins activate IRE1α through oligomerization and auto-transphosphorylation promoting RNAase activity to cleave XBP1 mRNA resulting in the expression of UPR effector XBP1s^11^. XBP1s, a transcription factor and specific marker of IRE1α mediated UPR activity, localizes to the nucleus to express genes involved in ER expansion, foldases, ERAD, secretion, and inflammation – where initial pro-survival UPR commits to cell death upon extended and unresolved ER stress^10, 11, 25, 27^. Our studies reveal that mitochondrial mutations sensitize cells to the activation of XBP1s and associated ER stress, and that tetracyclines can suppress its activation (**Fig. 3**). Tetracyclines were able to suppress ER stress chaperone BiP and also the phosphorylation of p38 downstream of IRE1α (**Fig. 3B**). The observation that ER chaperone capacity (BiP) is reduced upon mitoribosome inhibition implied that ER protein loading (ex. translocation) may be reduced under these conditions. Consistent with this, mitochondrial mutant cells exhibited strong increases in the loading of ER-protein cathepsin-C in isolated microsome fractions when compared to WT, and this loading was attenuated with tetracyclines (**Fig. 4**). Additionally, mitochondrial mutants were more dependent on ERAD for survival, where maladaptive protein loading may require this mechanism of protein homeostasis for survival in stressed conditions. This suggests that the activation of XBP1s in mitochondrial mutants may be due to increased ER protein loading, where inhibition of mitochondrial translation with tetracyclines, through an unknown mechanism, is attenuating ER protein translocation and accompanied ER stress. Activated UPR and SRP mediated co-translational translocation have been recently observed in patient fibroblasts with aberrant complex IV assembly^33^, substantiating our observations and highlighting a maladaptive response to mitochondrial mutations.

At ER-mitochondrial contacts, IRE1α oligomerization and activity is regulated by outer mitochondrial membrane E3-ubiquitin ligase MARCH5^25^. Reminiscent of tetracyclines, MARCH5-mediated ubiquitylation of IRE1α attenuates high order oligomerization, controlling the extent of UPR and cell fate. We illustrate that tetracyclines promote survival in MD cells independent of MARCH5 (**Fig. S2**). Mitochondrial stress, including translational stalling, can be communicated to the cytosol through the integrated stress response (ISR)^34, 35^. DELE1, a mitochondrial protein, is cleaved by protease OMA1 under mitochondrial stress conditions, where this DELE1 short fragment localizes to the cytosol, binds to activate the heme regulated kinase (HRI) phosphorylating eIF2α (p-eif2α) activating the ISR and expression of activating transcription factor 4 (ATF4)^34, 35^. To test if DELE1 is involved in the doxycycline survival mechanisms, we similarly used CRISPR Cas9 to KO DELE1 and score survival under galactose conditions (**Fig. S3**). These studies illustrated that doxycycline at higher concentrations (100 µM) acutely induced p-eif2α and ATF4 in a DELE1 dependent fashion, however these responses were not sufficient to reverse doxycycline induced survival (**Fig. S3**). This is consistent with our previous findings^7^ where knockdown of ATF4 with RNAi and inhibition of p-eif2α with ISRIB did not reverse doxycycline survival, disqualifying ATF4 and most likely other upstream ISR- transducers in the survival mechanism. Thus, we illustrate that tetracyclines promote survival independent of known mitochondrial proteins that are annotated to mediate stress responses: UPR targeting MARCH5 and ISR mediating DELE1, indicating a novel regulation of IRE1α originating at the mitochondria.

The maintenance of the mitochondrial proteome requires an intricate balance between the import of nuclear encoded proteins synthesized on cytosolic ribosomes, and the expression of mitochondrial encoded proteins from mitoribosomes^4^. Encoding genes for rRNA and tRNA, mtDNA also encodes 13 hydrophobic protein subunits of the electron transport chain complexes^6, 36^. Due to their hydrophobic nature, mitochondrial encoded nascent proteins are efficiently inserted into the inner mitochondrial membrane, where the mitoribosome LSU is tethered to insertase OXA1L, and specific chaperones direct ETC complex biogenesis with mitochondrial-derived subunits^36^. Elongational stalling of mitoribosomes during this process activates mitochondrial quality control factors^15^ where we illustrate that tetracyclines mediate a splitting and recruitment of MALSU1 to the mtLSU (**Fig. 2**) which is required for the alleviation of ER stress and cell survival (**Fig. 1, 3, & 4**). In this regard, we claim that general suppression of mitochondrial translation is not sufficient to promote survival of mitochondrial mutants, but rather, the action of tetracyclines in splitting the mitoribosome is required for the survival mechanism. These claims are substantiated with the observation that peptide deformylase inhibitor actinonin, a non-mitoribosome-targeting translation inhibitor, also did not promote survival (**Fig. 1A**). These mechanistic insights can be expanded to other antibiotics that target tRNA accommodation phase of mitoribosome elongation including pentamidine^19^, or mtLSU peptidyl transferase targeting pleuromutilins^13, 20^; compounds we identified and validated to rescue mitochondrial mutant cell death. Our data indicates that MALSU1-bound and elongating ribosomes constitute a signaling platform that communicates translation states in the mitochondria to the ER to resolve maladaptive protein loading. There is recent evidence of coordinate regulation between mitochondrial and cytosolic ribosomes^37, 38^ and thus, attenuation of mitochondrial translation by tetracyclines may induce a specific and currently unknown signal that reduces cytosolic protein loading in the ER.

## Materials and Methods

### Cell lines, treatments, and culture conditions

ND1 and WT cybrid cells previously utilized by us^5^ were generated by R. Vogel and J. Smeitink (Radboud University Medical Centre) using patient fibroblast mitochondria carrying either WT or the A3796G (ND1) mutation. Cybrid cells were cultured at 37⁰C and 5% CO2 in DMEM high glucose (25mM, Gibco) medium supplemented with sodium pyruvate (1mM), fetal bovine serum (10%), and penicillin-streptomycin (100 U/mL). For glucose deprivation experiments, DMEM no-glucose (Gibco, 11966-025) was supplemented with equimolar (25mM) galactose, sodium pyruvate (1mM), fetal bovine serum (10%), and penicillin-streptomycin (100 U/mL). Expi293F cells (Gibco) were cultured in suspension in Expi293 expression medium (A14351-01) without antibiotics using the manufactures protocol^39^. Cells were maintained at a density of 0.5 – 5x10^6^ cells/ml and cell viability (trypan blue) was assessed upon subculture to maintain viability >95%.

### Antibodies and reagents

Reagents used in this study include: Doxycycline hyclate (Sigma), actinonin (Sigma), eeyarestatin I (Sigma), 4µ8C (Sigma), n-dodecyl β-D-maltoside (β-DDM, Anatrace), cardiolipin (18:1, Avanti Polar Lipids), β, γ-methyleneguanosine 5’-triphosphate sodium salt (GMPPCP, Sigma).

Antibodies used in this study include: C12orf65 (mtRF-R, abcam, ab122448, 1:1000), C6orf203 (MTRES1, abcam, ab151066, 1:1000), C7orf30 (MALSU1, Proteintech, 22838- 1-AP, 1:1000), MRPL4 (Proteintech, 27484-1-AP, 1:1000), MRPS16 (Proteintech, 16735-1-AP, 1:1000), LRPPRC (Proteintech, 21175-1-AP, 1:1000), TOM70 (Santa Cruz, sc390545, 1:1000), IRE1α (Cell Signaling, 14C10, 1:1000), β-tubulin, Histone-H3 (abcam, ab52946, 1:1000), VDAC (Santa Cruz, sc390996, 1:1000), XBP1s (Cell Signaling, D2C1F, 1:1000), PARP (Cell Signaling, 46D11, 1:1000), BiP (Cell Signaling, C50B12, 1:1000), P- p38 (T180/Y182, Cell Signaling, D3F9, 1:1000), p38 (total, Cell Signaling, 1:1000), pan – OXPHOS (ATP5A1, UQCRC2, MTCO1, SDHB, NDUFB8, abcam, STN-19467, 1:500), Cathepsin-C (R&D Systems, AF1071, 1:1000), SRP14 (Proteintech, 11528-1-AP,1:1000), Calnexin (Cell Signaling, C5C9, 1:1000), MARCH5 (abcam, ab185054, 1:1000), HRI (EIF2AK1, abcam, 1:1000), p-eif2α (S51, Cell Signaling, 119A11, 1:1000), eif2α (total, Cell Signaling, 1:1000), ATF4 (Cell Signaling, D4B8, 1:1000).

### Cell survival assays

Survival assays of mitochondrial mutant ND1 cybrid cells were carried out as previously described by us^7^. Cells were seeded at 1.0x10^5^ cells/well in 6-well plates in glucose-deprivation media. Doxycycline, actinonin, or eeyarestatin I (ESI) were added at indicated concentrations at the time of seeding. Cells were then incubated for 96 hours at which time the surviving cells were counted via trypan-blue exclusion method. Selected experiments (**Fig. 3A**, **Fig 4A-B**) were performed in 24 well plate format where cell survival was assayed in response to time-course or dose responses, respectively. In time course experiments, ND1 cells were seeded in glucose deprivation media at 2.5 x10^4^ cells/well and doxycycline was added at the time of seeding. Cells were incubated for the indicated timepoints at which media was aspirated and cells were rinsed with PBS and air-dried to fix overnight. Cells were then stained with sulforhodamine B (SRB) and cell survival was scored as a function of total cellular protein as previously described^40^. ESI dose response curves were generated using SRB by seeding ND1 or WT cells with or without glucose at varying concentrations. Cells were then incubated for 48 hours after which media was aspirated and analyzed for cell survival via SRB as indicated above. Dose-response curves were generated and IC50 values were calculated using GraphPad prism software and illustrate the concentration at which 50% of cells survive when compared to non-treated control cells in each respective media condition (glucose or galactose).

### CRISPR-Cas9 gene editing

CRISPR-Cas9 gene editing experiments were performed using the GeCKO^41^ protocol. sgRNA guides were cloned into lentiCRISPR-V2 plasmids containing the puromycin resistance vector (Addgene, 98290), transformed into Stbl3 competent E. coli cells, and purified using standard methods. HEK293T cells were then seeded in 6-well dishes (6x10^5^ cells/well) and reverse transfected using Polyfect (Qiagen, 301105) following the manufacturers protocol. Transfection was carried out using 880 ng of lentiCRISPR-V2 with respective target- or non-targeting control sgRNA, 600 ng psPAX2 (Addgene, 12260), and 300 ng pMD2 (Addgene, 12259). Following 24 hr transfection, media was replaced with DMEM and virus was generated for an additional 48 hr. Lentiviral medium was then harvested, filtered (0.45 µm), and added with polybrene (conc, Cat. No.) onto ND1 cells that had been seeded and adhered to 6 well dishes (3x10^5^ cells/well) overnight. 24 hr after transduction, cells were split into 10 cm dishes and puromycin (0.5 µg/ml) was added and cells were selected for 7-10 days before used for experiments. The following sgRNA sequences were used: mtRF-R: 5’-ACCTTTACAACGATGCCTGA-3’. MTRES1: 5’- ATGATGTTGTCCTGAAGACG-3’. MALSU1: sg1 5’- CAGGCCGCGGACAAAGTTTG-3’; sg2 5’- CCCACTAATGTGGCGCAGGG-3’; sg3 5’- GCGGAGGGGACGGTCAACGA-3’. DELE1: 5’-TGATAATAAAGGACCGCCTG-3’. MARCH5: 5’- CCAGGCCTGTCTACAACGCT-3’.

### Dependency Map proteomics gene-ontology analysis

Proteomic correlation of doxycycline was scored and ranked by p-value (DepMap, PRISM repurposing screen^21, 22^), where proteins enriched in cells surviving under doxycycline treatment were identified (p<0.0001). Within this statistically significant subset of proteins, positive correlates were further analyzed for gene-ontology (GO) enrichment against the entire proteome using DAVID functional annotation bioinformatics (https://david.ncifcrf.gov/)42. GO-terms and their respective calculated fold-enrichment were ranked and plotted. High fold-enrichment represented a high number of proteins within a given GO-term that were enriched in doxycycline treated cells.

### Subcellular fractionations

Subcellular fractionations were carried out using differential centrifugation as previously described^43^. ND1 or WT cybrid cells were seeded in 2x15 cm dishes in glucose media and allowed to reach 80% confluence. Cells were then rinsed with PBS and galactose media was then added with doxycycline at indicated concentrations and were incubated for 48 hours. Cells were then harvested by scraping and pelleted at 300xG for 5 min at 4⁰C. Cells were resuspended in PBS, pelleted, and then resuspended in 5 ml of cold isolation buffer (IB, 225 mM mannitol, 75 mM sucrose, 0.1 mM EGTA, 30 mM HEPES-HCl pH 7.4). Cells were than dounce-homogenized (∼60 up-down strokes) on ice and centrifuged at 1000xG at 4⁰C for 5 min. Supernatant containing mitochondria, ER (microsomes), lysosomes, and cytosol, was then decanted into an additional 5 ml of IB and centrifuged at 10,000xG at 4⁰C for 10 min to obtain crude mitochondria (mitochondria/ER) in the pellet. Supernatant containing cytosol, microsomes, and lysosomes was decanted and kept on ice and mitochondria/ER fractions were resuspended in mitochondria resuspension buffer (MRB, 250 mM mannitol, 5 mM HEPES pH 7.4, and 0.5 mM EGTA). MRB was centrifuged at 10,000xG at 4⁰C for 10 min, supernatant was aspirated and crude mitochondria were snap frozen in liquid nitrogen and stored at -80⁰C. Lysosomes were then separated from microsomes and cytosol by centrifugation (20,000xG at 4⁰C for 30 min), pelleted and snap frozen. Microsomes in the supernatant were isolated by ultra-centrifugation (100,000xG for 60 min at 4⁰C) and resuspended in IB with protease inhibitors and snap frozen in liquid nitrogen and kept at -80⁰C. An aliquot of the resulting final supernatant containing cytosol was also snap frozen in liquid nitrogen and kept at -80⁰C.

### Western blotting and Blue-Native PAGE

SDS- and Native PAGE were carried out using standard protocols. For SDS-PAGE, cells were and resuspended in RIPA buffer (Cell Signaling, 9806) with protease and phosphatase inhibitors at 4⁰C and protein content was determined using standard BCA colorimetric protocols (Pierce 23228). Microsomes in IB and crude mitochondria in MRB (from subcellular fractionation) were directly quantified and solubilized with Laemmli SDS-sample buffer (Bio-Rad, 1610747). 10-20µg of protein from respective samples were run on 4-12% SDS-PAGE gels (Invitrogen, NP0336BOX). Proteins were then transferred to PVDF membranes and probed with specific antibodies. Native-PAGE analysis was carried out as previously described^24^ with some modifications. Mitochondria/ER (100 µg) were solubilized in 50 µl of Native Page Buffer (Invitrogen, 2360376) supplemented with 1.5% digitonin and protease inhibitors. Cells were then incubated at 4⁰C for 10 min to complete lysis, and centrifuged at 18,500xG for 20 min (4⁰C) to remove insoluble material. Supernatant was then carefully removed and resuspended in 0.5% G-250 (Invitrogen, BN20041). Samples (10 µg) were then added to each well and protein complexes were separated on 3-12% NativePAGE gels (Thermo Fisher Scientific, BN1003BOX). Proteins were transferred to PVDF membranes and probed with specific antibodies. Buffers used for NativePAGE include: NativePAGE Running Buffer (20X; Thermo Fisher Scientific, BN2001) and NativePAGE Cathode Buffer Additive (20X; Thermo Fisher Scientific, BN2002).

### Isolation of mitoribosomes

Mitoribosomes were isolated following a procedure similar to previously described^15^. Expi293F cells (750 ml, 6x10^6^ cells/ml) treated with doxycycline (1 µM) or vehicle (DMSO) for 48 hours were harvested by centrifugation (300xG, 5 min, 4⁰C). Cells were resuspended in cold PBS and re-pelleted. Cells were then resuspended in mitochondrial isolation buffer (MIB, 50 mM HEPES-KOH pH 7.5, 10 mM KCl, 1.5 mM MgCl2, 1 mM EDTA, 1 mM EGTA, 1 mM DTT, and protease inhibitors) at a ratio of 6 ml MIB per 1 g of pellet. Cells were left to swell at 4⁰C for 10 min, after which were brought to isotonic concentrations of sucrose and mannitol by adding calculated volumes of 4X sucrose mannitol buffer (SM4, 280 mM sucrose, 840 mM mannitol, 50 mM HEPES-KOH pH 7.5, 10 mM KCl, 1.5 mM MgCl2, 1mM EDTA, 1 mM EGTA, 1 mM DTT, and protease inhibitors). The resulting solution MIB + SM4 (MIBSM) will be used in following steps. Cells were then dounce homogenized (∼60 up- down-passes) and cellular debris and nuclei were centrifuged (800xG, 4⁰C, 15min). Supernatant-1 was filtered and pellet was resuspended in fresh MIBSM for 15 passes. Resulting solution was again centrifuged and supernatant- 2 was combined through a cheesecloth with supernatant-1. This solution was clarified by centrifugation a (1,000xG at 4⁰C for 15 min) and resulting supernatant was collected. Crude mitochondrial pellet was then obtained by centrifugation (10,000xG at 4⁰C for 15 min), and was resuspended in MIBSM buffer with DNAase (0.1U/ml) and left rocking at 4⁰C for 30 min. Mitochondria were pelleted (10,000xG at 4⁰C for 15 min), resuspended in 1 ml of SEM buffer (250 mM sucrose, 20 mM HEPES-KOH pH 7.5, 1 mM EDTA) and loaded onto a discontinuous sucrose gradient (15%-60%) constructed as previously described^15^. Mitochondria were obtained, snap frozen in liquid nitrogen, and kept at -80⁰C.

To obtain mitoribosomes, mitochondria were thawed at 4⁰C and combined with equal parts of lysis buffer (25 mM HEPES-KOH pH 7.4, 150 mM KCl, 50 mM MgOAc, 1.5% β-DDM, 0.15 mg cardiolipin, 0.5mM GMPPCP, 2 mM DTT, and protease inhibitors). Combined solution was mixed and gently homogenized with a dounce homogenizer and was left to rock at 4⁰C for 30 min. Lysed material was centrifuged at 20,000xG for 30 min to clarify, and supernatant containing mitoribosomes was carefully decanted. Lysis solution (2 ml) was added on top of a sucrose cushion (1 ml, 1M sucrose (34%), 20 mM HEPES-KOH pH 7.4, 100 mM KCl, 20 mM MgOAc, 0.6% β-DDM, 0.06 mg/ml cardiolipin, 0.25 mM GMPPCP, 2 mM DTT) in SW 55 Ti tubes. Samples were centrifuged (231, 550xG at 4⁰C for 1 hr) to obtain crude mitoribosome pool. Mitoribosomes were reconstituted in 100 µl of resuspension buffer (20 mM HEPES-KOH pH 7.4, 100 mM KCl, 20 mM MgOAc, 0.3% β-DDM, 0.03 mg/ml cardiolipin, 0.25 mm GMPPCP, 2 mM DTT) and loaded on top of a 13- 30% linear sucrose gradient^15^ in SW 55 Ti tubes. Samples were centrifuged (213,626xG at 4⁰C for 90 min), and entire volume was fractionated into 13 equal parts. Samples were snap frozen and kept for western blotting analysis. Inputs represent crude mitoribosome pool. ND1 mitoribosomes were analyzed in a similar fashion, but crude mitochondria (see *subcellular fractionation*) were directly lysed and subjected to linear sucrose gradient fractionation without purification of mitoribosome pool. Inputs represent crude mitochondria in these experiments.

### Quantitative PCR

Cells were seeded (1.0x10^5^ cells/well) in 6 well dishes and treated with the indicated compounds for 48 hours under glucose starvation conditions. RNA was then isolated using Trizol (Invitrogen, 15596-026) and a Zymo-Spin Direct-zol RNA Kit (Zymo Research, R2050) following the manufactures instructions. 1 μg of RNA was then used to reverse transcribe cDNA using a High-Capacity cDNA Reverse Transcription Kit (Applied Biosystems, 4368813) following. cDNA was then mixed with SYBR Green qPCR master mix (Applied Biosystems, 4309155) and amplified on a CFX 384 Real-Time system (Bio- Rad) for analysis. Primers used for this study include Human: MALSU1 F- CTGCTCGCCCCACTAATG, R-ACTTGGGACCAGTATGATCTGC.

### Cycloheximide chase experiments

ND1 or WT cybrid cells were seeded in glucose in 2 x 15 cm^2^ dishes per condition and allowed to reach 90% confluence. Cells were pulsed with cycloheximide (200 µg/ml) for 0, 3, and 6 hours after which they were harvested and snap frozen in liquid nitrogen. Whole cells and microsomes were then isolated (see *subcellular fractionations*) for western blot analysis.

### Animal experiments

Animal studies were performed as previously reported by us^7^. C57BL/6J NDUFS4-/- or WT mice supplemented with doxycycline or normal chow diets were euthanized at ∼p55 and brains were immediately harvested and snap frozen in liquid nitrogen and kept at - 80⁰C. Brain tissue was then pulverized with mortar and pestle until a fine powder was obtained. Tissue was then resuspended in RIPA buffer for western blot analysis.

### Ethical considerations

Animal studies were performed in compliance with the Institutional Animal Care and Use Committee protocol approved by the Beth Israel Deaconess Medical Center Animal Facility.

### Statistics

Replicates presented are derived from independent biological samples. Statistical analysis between two samples was done using student t-test, and greater than two groups using one- or two-way ANOVA with multiple comparisons where appropriate. Statistical significance was determined with p-value <0.05*, <0.01**, <0.001***, and <0.0001**** as noted.

## Acknowledgments

We thank the entire Puigserver laboratory for helpful discussions and insights for the duration of the project. We also would like to extend our gratitude to the department of cancer biology at Dana Farber and department of cell biology at Harvard Medical School for their support. This work was funded by the NIH grants, R56 AG074527/AG/NIA and RO1 CA181217/CA/NCI (to P.P), T32 CA236754-02 (to C.T.R.), F32 GM125243-01A1 NIGMS (to C.F.B.), and F30 DE028206-01A1 NIDCR (to E.A.P.).

## Supplementary Figures

**Figure S1.**
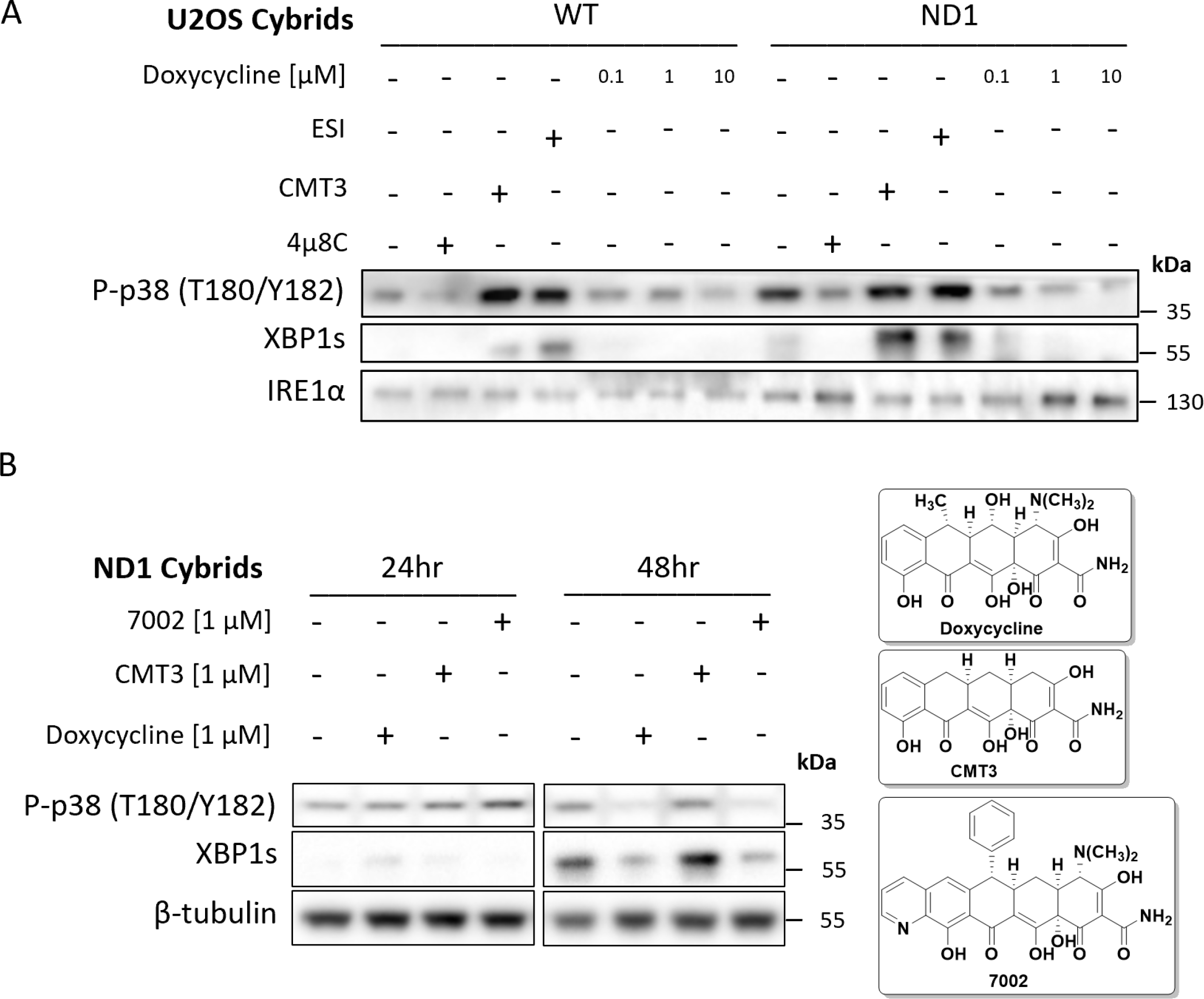
The activation of p38 MAPK is downstream of IRE1α and can be modulated by ERAD and mitochondrial translation. (**A**) WT and ND1 cells were cultured in galactose media for 48 hours and analyzed for the activation of MAPK and UPR signaling as evidenced by phosphorylation of p38 (P-p38) and expression of XBP1s, respectively. Pharmacological inhibition of IRE1α with 4µ8C (20 µM) and p97 inhibitor ESI (20 µM) indicate P-p38 and XBP1s are downstream of IRE1α and can be controlled by ERAD. CMT3 (1 µM) does not affect P-p38 MAPK or XBP1s levels when compared to doxycycline (1 µM) indicating inhibition of mitochondrial translation, not off target effects of tetracyclines, modulates UPR and MAPK signaling. (**B**) UPR and MAPK signaling are activated after 48 hours of galactose stress in ND1 cybrids, where doxycycline (1 µM) and chemically distinct 7002 (1 µM), but not CMT3 (1 µM) attenuate these responses.

**Figure S2.**
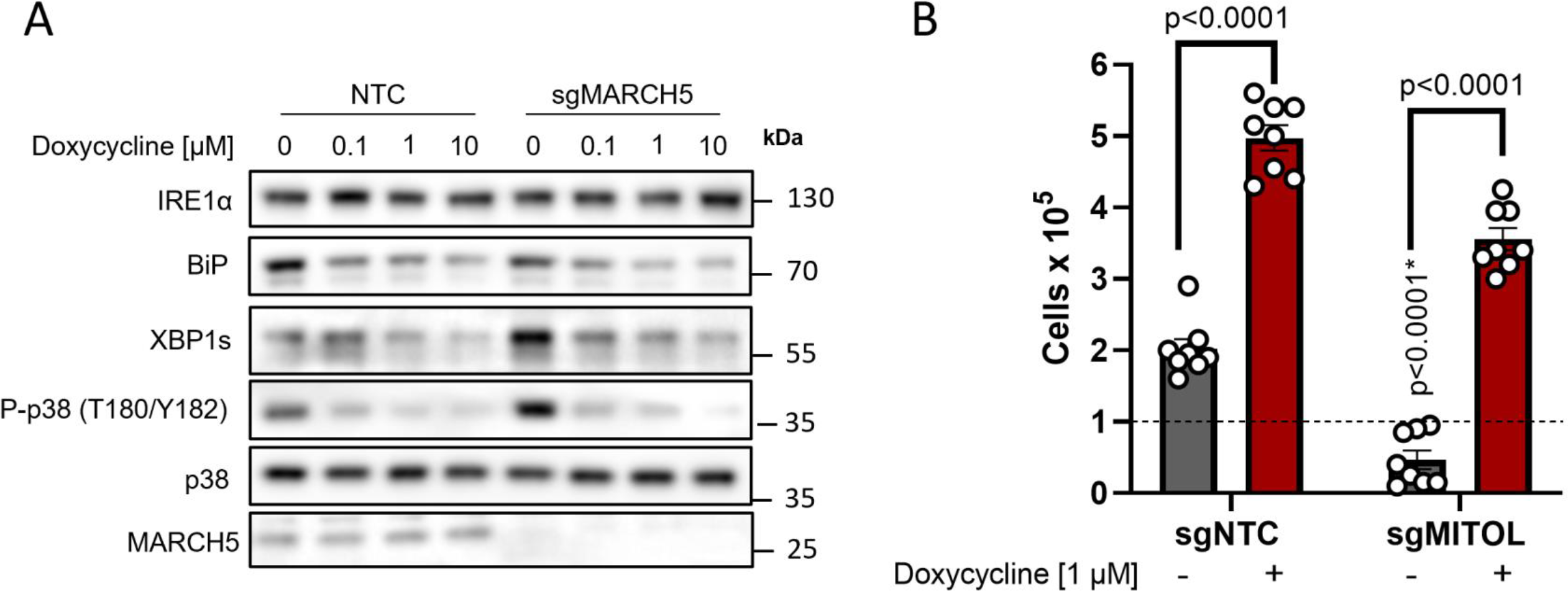
Outer-mitochondrial membrane E3-ubiquitin ligase MARCH5 is not required for tetracyclines pro-survival mechanism in mitochondrial mutant cells. (**A)** Doxycycline dose-dependently inhibits UPR and MAPK signaling in ND1 non-target control (NTC) cells and cells depleted of MARCH5 under galactose conditions. Note basal levels of XBP1s and P-p38 are increased in sgMARCH5 cells, indicating enhanced sensitivity to ER stress under these conditions. (**B**) ND1 cybrids cultured in galactose conditions for 4 days can be rescued by doxycycline independent of MARCH5 expression status (n=8 independent biological replicates across N=3 independent experiments).

**Figure S3.**
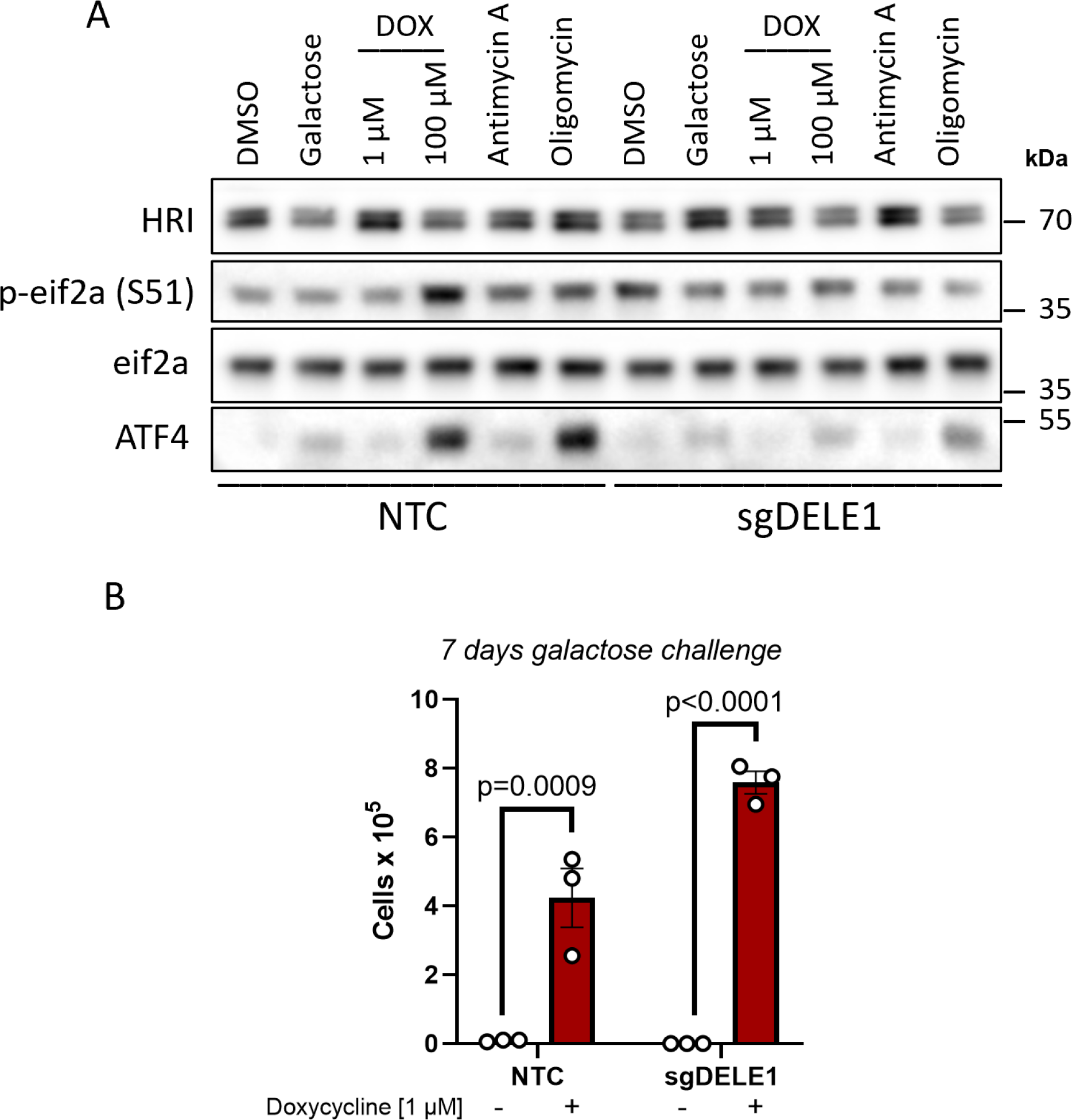
Activation of the integrated stress response with mitochondrial DELE1 is not required for tetracyclines to promote survival in mitochondrial mutant cells. (**A**) ND1 cybrid cells treated with mitochondrial targeting agents (6 hours) activate the integrated stress response as indicated by increases in P-eif2α and ATF4. Note changes in P-eif2α and ATF4 seen in NTC cells are suppressed in cells lacking DELE1 (sgDELE1). Cells were cultured under glucose conditions except for where noted (galactose). Doxycycline (1 µM, 100 µM), antimycin A (40 nM), and Oligomycin (1.25 ng/µl). (**B**) ND1 cells deprived of glucose for 7 days can be rescued by doxycycline (1 µM) independent of DELE1 expression status (n=3 biological replicates).

